# On the Origins of the Cerebral IVIM Signal

**DOI:** 10.1101/158014

**Authors:** Hannah V. Hare, Robert Frost, James A. Meakin, Daniel P. Bulte

## Abstract

**Purpose:** Intravoxel incoherent motion (IVIM) has been proposed as a means of non-invasive MRI measurement of perfusion parameters such as blood flow and blood volume. Its main competitor in the brain is arterial spin labelling (ASL). In theory, IVIM should not suffer from some of the same limitations as ASL such as poor signal in white matter, and assumptions about arterial arrival times that may be violated in the presence of pathology.

**Methods:** In this study we aimed to test IVIM as a viable alternative to ASL for quantitative imaging of perfusion parameters in the brain. First, a direct comparison was performed between IVIM and multi-post label delay pseudo-continuous ASL; second, IVIM images were acquired with and without nulling cerebrospinal fluid; and finally, ultra-high resolution IVIM was performed to minimise partial voluming.

**Results:** In all three tests, IVIM failed to disprove the null hypothesis, strongly suggesting that, at least within the brain, the technique does not measure perfusion parameters as proposed.

**Conclusion:** Furthermore, the results obtained suggest that the contrast visible in IVIM-derived images is primarily sensitive to cerebrospinal fluid, and not the microvascular blood compartment.

## Introduction

Intravoxel incoherent motion (IVIM) is a unique magnetic resonance imaging (MRI) approach to measuring blood volume and perfusion in the body (1). As its name suggests, it relies only on the motion of water molecules within a single voxel, and it should thus be capable of making equally valid measurements throughout the entire vascular network, including within white matter in the brain. This benefit may outweigh the method’s relative disadvantages compared to arterial spin labelling (ASL) (functional imaging would be difficult as a full dataset is required to derive a single perfusion map), whilst still allowing for non-invasive imaging of cerebral perfusion, especially in the presence of very slow-moving blood. The aim of this work was to assess the viability of using IVIM to measure resting cerebral blood volume and perfusion.

ASL is capable of non-invasively measuring cerebral perfusion using MRI in a clinically feasible imaging time. Although it suffers from poor signal-to-noise (SNR), recent advances in the suppression of background signal and Bayesian analysis approaches have enabled a transition from research into clinical practice (2), and several institutions now include ASL as a standard sequence in their protocols. However, the ASL signal is fundamentally limited by the T_1_ decay of the blood, so that even very long post-label delay times are not able to reliably quantify very slow-moving blood. White matter perfusion is notoriously difficult to measure using ASL (3); similarly, it is very challenging to obtain perfusion values in areas of cerebral occlusions or in the presence of collateral flow, which are typically identified on ASL images by a lack of blood signal. This creates particular challenges for other MRI techniques which rely on ASL data for perfusion information, such as calibrated MRI. To date, calibrated MRI techniques have been applied to investigate various aspects of the healthy human brain, both at rest and in response to a range of functional stimuli, but they have rarely been applied in cases of pathology, even within a research setting (4–6). At the time of writing only a single study had applied the method in patients. De Vis et al. reported artefacts arising from delayed arrival of arterial blood in 3 out of 11 patients scanned with internal carotid artery occlusions, leading to unreliable estimates of oxygen extraction and consumption (7). Thus the areas likely to be of greatest interest in the monitoring of oxygen metabolism are simultaneously the areas of least confidence for ASL perfusion measures.

Velocity-selective ASL was developed to overcome this limitation by creating a tagging scheme sensitive to the change in velocity of arterial water molecules rather than their position (8). However, this technique suffers from even lower SNR than other flavours of ASL, at least for normal arterial arrival times.

Intravoxel incoherent motion was first developed by Le Bihan in the 1980s (1), and is based on the premise that both diffusion and microvascular perfusion result in motion of water molecules in all directions (at the macroscopic scale of a voxel), but at different temporal scales, with water in the capillaries moving more quickly than water in intra-cellular space. This leads to a biexponential model of diffusion signal as a function of b-value, with very low b-values (less than ~100 s/mm^2^) exhibiting a greater rate of signal decay due to capillary flow. The MR signal will be attenuated by both diffusion and perfusion according to

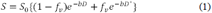

In addition to the classical diffusion coefficient (*D*), a ‘pseudo-diffusion coefficient’ *D*^*^ is introduced, along with the fraction of fast-diffusing spins, *f_v_*, which is usually interpreted as the capillary blood volume fraction in each voxel. The product *f_v_D*^*^ is then a measure of perfusion, in units of mm^2^/s, which may be converted to traditional CBF units of ml blood/100 g tissue/minute via the following equation (9):

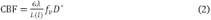

where λ is the fraction of MR-visible water, *L* is the total length of the capillary network and 〈*l*〉 is the mean capillary length. Capillary lengths cannot be measured non-invasively; they have been shown to vary with age (10) and region (11) within subcortical structures, but appear to be remarkably constant across the cortex (10,12). However, they may be expected to vary significantly in cases of pathology. For this reason *f_v_D*^*^ values are not usually converted to traditional perfusion units. The term ‘perfusion’ will be used throughout this manuscript with the understanding that it refers to the pattern of blood flow through the capillaries, and not specifically to the delivery of oxygen or nutrients to the tissue. This is true for both ASL and IVIM imaging.

IVIM has been used to study the liver (13–17), kidney (18–20) and pancreas (21–23) without the need for an injected tracer, which can be particularly valuable in cases where contrast agents are contraindicated. However, its application within the brain has been a subject of controversy for many years (9,24,25). Results can vary significantly depending on the specific fitting algorithm that is used (26), as well as on experimental parameters such as TE (23) and the range of b-values measured (27,28). Concerns have also been raised about biases introduced by fitting low SNR data (29,30), and contaminating effects of cerebrospinal fluid (CSF) due to bulk motion during the cardiac cycle or simple partial voluming (31).

Despite these complications, IVIM continues to be applied in the brain as well as numerous abdominal organs. IVIM-derived parameters have shown promise in the characterisation and monitoring of head and neck tumours (32–36). Going one step further, Federau et al. have reported quantitative changes in IVIM perfusion during hypercapnia (37) and visual stimulation (38), despite earlier work failing to achieve this in animal models (39).

The aim of this work was to investigate IVIM as a potential method for acquiring reliable, quantitative blood volume and/or perfusion information in the healthy brain. First, a direct comparison was performed between IVIM and ASL, a method capable of quantifying blood flow within grey matter which has been repeatedly validated against PET (40–42). Second, the impact of CSF on the IVIM signal was assessed by application of a CSF-nulled diffusion sequence. Finally, high-resolution IVIM data were acquired to isolate and compare signals arising from grey matter, white matter and CSF.

## Methods

### Experiment 1

10 subjects were scanned on a 3 T Siemens Verio scanner with a 32-channel head coil under FMRIB’s developmental ethics agreement. A standard 6:28-minute multi-post label delay pseudo-continuous ASL (PCASL) scan was run (6 delay times ranging 250–1500 ms) with 15 slices and voxel dimensions 3.4×3.4×6 mm (43). Head and body coil calibration images and field maps were also acquired. All PCASL data were pre-processed using FSL tools for motion and field map correction, and analysed using the BASIL toolbox (44).

For the IVIM, a standard diffusion sequence was run with b-values of 0, 10, 20, 40, 80, 110, 140, 170, 200, 300, 400, 500, 600, 700, 800 and 900 s/m^2^, acquired in three orthogonal directions with a twice-refocused spin echo (45) and an EPI readout. Simulations have suggested that a minimum of 10 b-values should be acquired in order provide sufficient points for reliable biexponential signal fitting (27), and the distribution used in the present study was chosen with reference to recent literature applying IVIM in the brain within a clinical setting (35). Other parameters included TE 84 s, TR 4 s, bandwidth 1086 Hz/pixel, echo spacing 0.99 ms, Fourier 6/8, GRAPPA off. Images were acquired at the same matrix size and resolution as PCASL data. In order to match the PCASL imaging time, 2 repetitions were acquired and averaged on the scanner, with an imaging time of 6:12 minutes. A b = 0 image with reversed phase-encoding was also acquired.

A 1 mm isotropic MPRAGE structural scan was acquired, which was segmented using FMRIB’s Automated Segmentation Tool (FAST) (46) and registered to both ASL and IVIM space using FLIRT (47). Grey matter (GM) and white matter (WM) masks were created from voxels with partial volume estimates of 50% or above.

### Experiment 2

One subject was re-scanned with an adapted protocol which was designed to identify signal arising from the CSF compartment. A non-selective bandwidth-modulated adiabatic selective saturation and inversion (BASSI) pulse was applied to null signal from spins with T_1_ = 3700 ms (inversion region thickness = 10,000 mm) (48), and only a single slice was acquired to ensure optimal CSF suppression. Imaging time and other parameters were identical to those set out in Experiment 1. For comparison, a second set of IVIM images was acquired with the BASSI voltage set to 0 for no CSF suppression. A PCASL scan was also performed, as described above.

Because of anticipated difficulties in registering a single slice to a structural image, a double inversion recovery sequence was run at the same resolution as the IVIM diffusion scans to obtain a suitable grey matter mask. Inversion times were set at 550 and 4150 ms prior to the excitation pulse. The resulting image was field map corrected using FUGUE (49) and manually thresholded to create a grey matter mask.

### Experiment 3

In order to analyse IVIM signals definitively originating only from a single tissue type (i.e. with minimal partial voluming), 6 subjects were scanned on a 3T Siemens Prisma with a high-resolution readout-segmented EPI (rs-EPI) diffusion-weighted sequence (50) using a 32-channel head coil. This allowed for a 1 mm isotropic resolution with minimal distortions, such that partial voluming could be avoided in a substantial subset of voxels. In order to maintain sufficient SNR the 6:14-minute protocol was repeated 6 times. Because of the potential for motion over this long scan time, the images were not averaged. The rs-EPI sequence was set up with TE 69 ms, TR 1.6 s, bandwidth 766 Hz/pixel, 5 readout segments, echo spacing 0.4 ms, GRAPPA factor 2. b-values were identical to those in Experiments 1 and 2, but were acquired with Stejskal-Tanner (single-refocused spin echo) encoding to minimise TE and maximise SNR (51).

An MPRAGE scan was performed at the same resolution and matrix size as the diffusion scan. This was segmented using FAST (46) to estimate the relative proportions of GM, WM and CSF in every voxel. Due to the increased spatial resolution and scan time masks could be created from voxels with 100% of the relevant tissue type.

### IVIM Fitting

IVIM data from Experiments 1 and 3 were first corrected for motion and eddy current distortions (52). The reversed phase-encoding b = 0 scan was used to correct for susceptibility-induced distortions in Experiment 1 using the TOPUP tool (53). Because the data acquired in Experiment 2 was for a single slice only, it was not possible to apply eddy corrections or TOPUP. Instead, a field map was used in conjunction with FUGUE (49) to reduce distortions near the sinuses.

A biexponential model was fitted to each voxel in two steps (26,37) using least squares fitting in MATLAB (MathWorks, Natick, MA, USA). This approach aims to mitigate the problem of overfitting, under the assumption that *D*^*^ is substantially larger than *D* and may be neglected for large b-values. First, a monoexponential model was fit to b-values greater than 200 to estimate *D*:

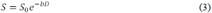

Then the full biexponential model was fit to all b-values, with fixed *D*, to estimate *f_v_* and *D*^*^:

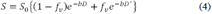

*S_0_* was unconstrained in both cases.

### Statistical Comparisons

For Experiment 1, GM masks were applied to ASL and IVIM images. Voxels with *f_v_* > 0.3 were excluded from masks as this is unphysiological, presumably as a result of CSF contamination (35). Correlations between ASL perfusion and IVIM *f_v_* and perfusion were hypothesised to be linear and were assessed by computing the Pearson product-moment correlation coefficients (r) and the corresponding p-values, where p < 0.05 was deemed to be significant.

In Experiment 2, IVIM analysis was performed both on a voxelwise and a region of interest (ROI) basis. For ROI fitting, the signal was averaged across all b-directions and all GM voxels in order to boost SNR.

Voxelwise fitting was performed on data from Experiment 3, and the IVIM-derived parameter maps were visually compared to structural images. Correlations between blood volume fraction *f_v_* and GM, WM and CSF fraction of partially volumed voxels were investigated by plotting 2D histograms. Box plots were created to compare the distributions of *f_v_* and perfusion in pure GM, WM and CSF.

## Results

### Experiment 1

Average grey matter results are presented in table 1. White matter perfusion cannot be reliably measured using ASL because of the long arterial arrival time (3); however, no such limitation applies to IVIM. *f_v_D*^*^ in white matter was found to be 0.81 ± 0.20 × 10^−3^ m^2^/s, leading to a GM/WM perfusion ratio of 1.87 ± 0.88 (ranging from 1.12 to 3.97 in individual subjects).

**Table 1:** Average grey matter values for each subject in Experiment 1. *f_v_D*^*^ is the IVIM perfusion parameter. ASL units are ml/100 g/min; *D*, *D*^*^ and *f_v_D*^*^ are all measured in m^2^/s.

| Subject | ASL<br>perfusion | $f_v$ | $D$<br>($\times 10^{-3}$) | $D^*$<br>($\times 10^{-3}$) | $f_v D^*$<br>($\times 10^{-3}$) |
| --- | --- | --- | --- | --- | --- |
| 1 | 43.6 | 0.11 | 0.93 | 10.40 | 0.96 |
| 2 | 40.8 | 0.11 | 0.92 | 12.71 | 1.56 |
| 3 | 59.0 | 0.11 | 0.92 | 9.00 | 0.93 |
| 4 | 54.7 | 0.10 | 0.84 | 12.09 | 1.22 |
| 5 | 55.2 | 0.12 | 0.92 | 15.44 | 2.54 |
| 6 | 46.0 | 0.13 | 0.94 | 10.17 | 1.27 |
| 7 | 44.7 | 0.10 | 0.95 | 11.52 | 1.25 |
| 8 | 41.3 | 0.11 | 0.88 | 14.79 | 1.94 |
| 9 | 51.4 | 0.10 | 0.89 | 9.46 | 1.02 |
| 10 | 65.2 | 0.12 | 0.92 | 12.09 | 1.74 |
| mean ± std | 50.2 ± 8.2 | 0.11 ± 0.01 | 0.91 ± 0.03 | 11.77 ± 2.14 | 1.44 ± 0.51 |

Figure 1 shows ASL and IVIM maps for a subset of slices acquired from a representative subject, along with the GM mask used for IVIM parameters. The GM masks for ASL are almost identical to those for IVIM, with small differences arising from slightly different distortions between the two readouts and any motion between the scans. Maps for ASL and IVIM perfusion look qualitatively similar, with the same features visible on both sets of images. Quantitatively however, no significant correlation was observed between ASL and IVIM perfusion (figure 2(a), r = 0.090, p = 0.805) or ASL and IVIM f_v_ (figure 2(b), r = 0.107, p = 0.768).

**Figure 1:**
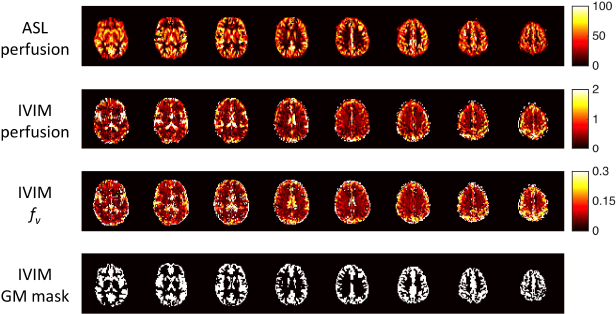
A subset of slices acquired from a single representative subject. Units are ml/100 g/min for ASL perfusion and 10^−3^ m^2^/s for IVIM perfusion. *f_v_* is the capillary blood volume fraction in each voxel.

**Figure 2:**
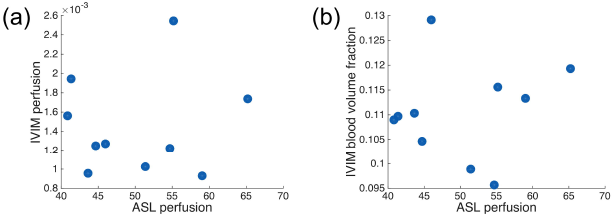
Plots comparing ASL perfusion with IVIM perfusion across the 10 subjects (a) and IVIM blood volume fraction (b) in grey matter. No significant correlation was observed in either case.

### Experiment 2

Figure 3 shows the results of the voxelwise fitting of both standard and CSF-suppressed IVIM sequences, along with an ASL perfusion map of the same slice and the GM mask created from a double inversion recovery sequence. CSF signal was successfully suppressed, as may be seen by comparing the b = 0 images in the top row. Both blood volume fraction and perfusion maps from IVIM fits bear some resemblance to the ASL map for the case of standard IVIM; however, this similarity disappears in the CSF-suppressed case.

**Figure 3:**
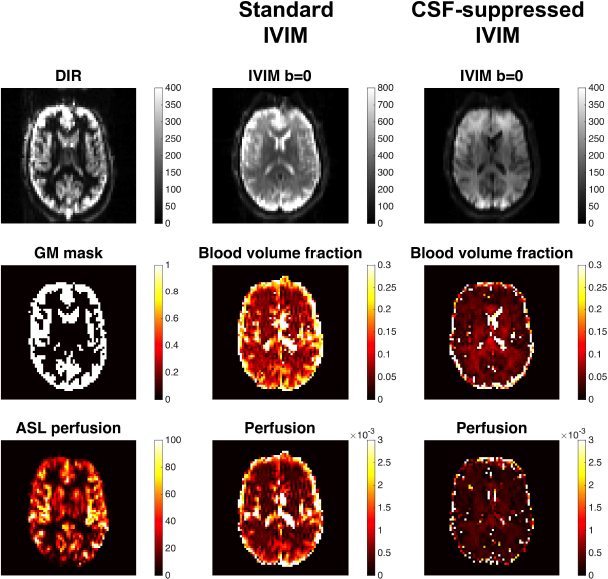
Single-slice IVIM results from Experiment 2, acquired with and without CSF-suppression. b = 0 images demonstrate that CSF was successfully suppressed by the sequence. Also shown is the double inversion recovery (DIR) image, which was thresholded to produce the grey matter (GM) mask. Blood volume fraction is *f_v_*, IVIM perfusion is *f_v_D*^*^ in units of m^2^/s. ASL perfusion has units of ml/100 g/min.

The ROI-averaged fits for standard and CSF-suppressed IVIM signals are shown in figure 4. For the standard case, the signal clearly exhibits biexponential behaviour. With CSF suppression the fits perform almost equally well; statistically the biexponential model produces a marginally higher R^2^, which appears to be driven by a closer fit for b < 40 s/m^2^, but visually there is little difference between the two fits.

**Figure 4:**
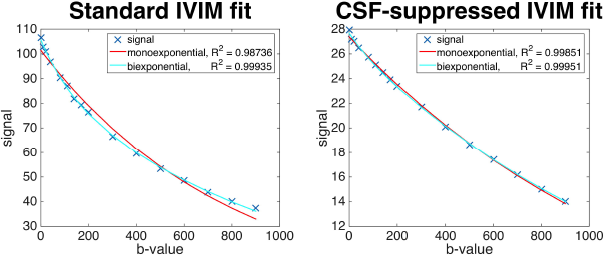
Plots showing the mono- and biexponential fits to IVIM signals from standard and CSF-suppressed data, after averaging over all grey matter voxels. The biexponential model is clearly a better fit for the standard IVIM data, whereas the monoexponential model appears sufficient for CSF-suppressed data. Note the different signal scales for the two cases.

### Experiment 3

Figure 5 shows maps of IVIM parameters from a single representative subject, alongside a structural image and the CSF partial volume estimate map derived from it. Areas with the highest IVIM blood volume fractions *f_v_* clearly correspond to those with the highest CSF fraction, and no obvious contrast between grey and white matter is seen in IVIM maps. All fits were done on a voxelwise basis with no smoothing or averaging applied. For reference, fits for 3 randomly selected voxels from the same subject, which contain 100% GM, WM and CSF respectively, are shown in figure 6.

**Figure 5:**
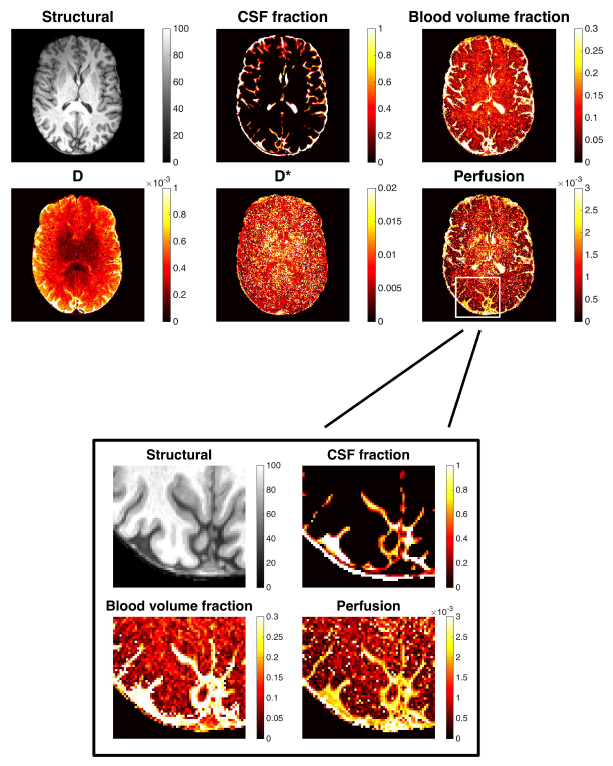
High-resolution IVIM data acquired with a rs-EPI diffusion sequence. Both blood volume fraction (*f_v_*) and perfusion (*f_v_D*^*^) are elevated in voxels containing high fractions of CSF, and no noticeable contrast is observed between grey and white matter.

**Figure 6:**
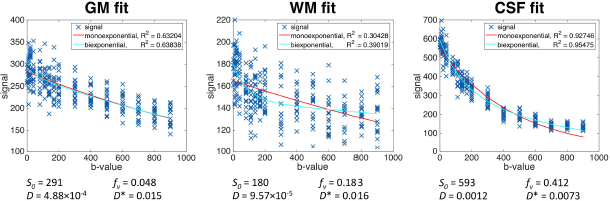
Plots of IVIM fits to single voxel data, taken from randomly selected pure tissue voxels in the slice shown in figure 5. Each b-value has 18 data points corresponding to 3 diffusion directions and 6 repeats. R^2^ values are highest in CSF voxels, presumably because of higher signal levels (see standard IVIM b = 0 image in figure 3 for an example of raw diffusion data). The monoexponential fits shown are for comparison only and were derived from all 16 b-values; they do not represent the fits performed to calculate D.

Structural images and blood volume fraction (*f_v_*) maps for all subjects are displayed in figure 7. Also shown are 2D histograms exploring the relationship between GM, WM and CSF partial volume estimates (pve) and IVIM blood volume fraction. No correlation is seen for GM or WM, but a positive trend is observed for CSF. Note that extreme values (100% pve voxels, *f_v_* = 1) have been excluded from these histograms. Figure 8 compares the distribution of *f_v_* and *f_v_D*^*^ in the form of box plots for a single subject, derived only from voxels with no partial voluming. These show considerably higher *f_v_* and *f_v_D*^*^ in CSF compared to GM or WM. Extreme outliers were observed for all tissue types and were excluded from the plots. *f_v_D*^*^ was significantly different in CSF compared to WM (p < 0.001 in all subjects) and GM (p < 0.05 in all subjects, and p < 0.001 in 4 of the 6). *f_v_D*^*^ in GM and WM differed significantly in only 3 subjects (p < 0.05), but in all 6 cases the median values were slightly higher in WM than in GM.

**Figure 7:**
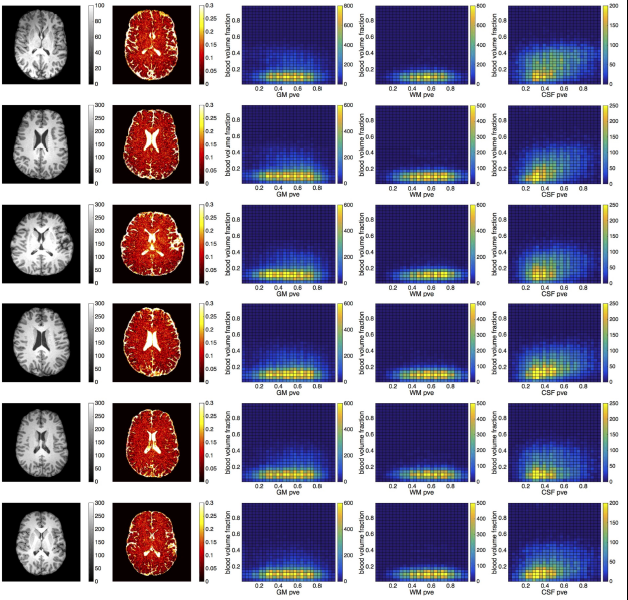
Results of Experiment 3 in all 6 subjects. Columns display one slice of the structural image (arbitrary units) and the blood volume fraction f_v_, followed by histograms correlating voxelwise GM, WM and CSF partial volume estimates (pve) against blood volume fraction. Voxels containing only a single tissue type are not included in these plots.

**Figure 8:**
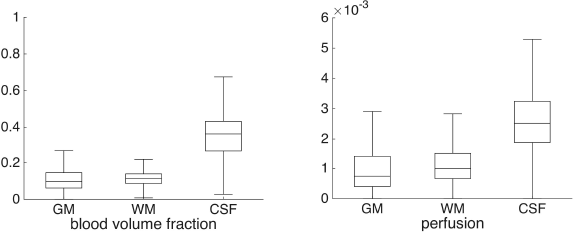
Box plots of the blood volume fraction (*f_v_*) and perfusion (*f_v_D*^*^) values observed in pure GM, WM and CSF voxels for a single representative subject.

## Discussion

IVIM claims to be a quantitative model (9), yet conclusive validation of the method within the brain is still lacking, with past studies reporting both positive (54) and negative (55) correlations between *f_v_* and DSC or ASL MRI methods, respectively. Although qualitative similarities were observed between IVIM parameters and ASL perfusion maps (figure 1), this study found no significant correlation between average GM values across a group of 10 subjects (figure 2).

At a low spatial resolution, contrast between grey and white matter is markedly reduced but still visible in IVIM parameters compared to ASL (figures 1 and 3). This is unsurprising given the limitations on arterial arrival time that ASL is subject to, meaning that WM perfusion will always be underestimated in ASL images. The mean ratio of GM/WM perfusion of 1.87 for IVIM compares well with the ratio of 2.07 observed in young healthy volunteers using PET (56), although the individual range in this study (1.12–3.97)was fairly large.

Using literature values for the mean capillary length <l> and the total length of the capillary network L it should be possible to convert *f_v_D*^*^ to traditional units of CBF using equation 1. For the human brain, values of λ = 0.78 [29], < *l* > = 53 μm (12) and *L* = 1 mm (57) would result in a CBF of 21.2 ml/100 g/min, somewhat lower than the ASL-derived 50.2 ml/100 g/min in the same cohort. As the definition of what exactly constitutes a capillary is not always clear, it is plausible that the scarce literature on human brain morphometry was based on a slightly different definition than that used in the derivation of the IVIM model.

It is possible that the lack of correlation shown in figure 2 could be caused by different vessel properties (e.g. average length of capillary segments) between individuals. Although the ability to compare absolute perfusion between subjects is very valuable, particularly in clinical cases of potential global hypo- or hyperperfusion, the scientific community is typically more concerned with generating spatial or functional contrast. Thus the question of greatest interest is whether the contrast observed in *f_v_* and *f_v_D*^*^ for a single subject is a true reflection of underlying blood volume or perfusion. Given past concerns over the contaminating effects of CSF signal (31), it is difficult to conclude whether the spatial similarity between ASL and IVIM images in Experiment 1 supports this hypothesis, especially as the majority of GM voxels will also include a sizeable fraction of CSF at this relatively low resolution.

Experiment 2 was designed to try to answer this question by acquiring diffusion data with and without CSF-suppression. At first glance, the results imply that the GM/WM contrast observed in IVIM-derived images is a direct result of CSF signal contamination. When CSF is successfully nulled, *f_v_* and *f_v_D*^*^ values are notably decreased in GM (figure 3), and a simple monoexponential model is adequate for fitting the data (figure 4). This is in agreement with past literature (31).

However, there are several limitations to the method used in Experiment 2. Only a single inversion preparation pulse was implemented; this was timed to maximally suppress signal from the CSF compartment, but inevitably will also have reduced signal from other tissue types. Previous simulation work suggests that when CSF is nulled following a single inversion, signal from arterial blood is reduced to 30% and GM signal to 40% of their respective equilibrium M_0_ values (58). Thus the CSF-suppressed data not only has lower SNR compared to the standard case, but also differing relative fractions of blood, GM and WM signal contributions. This makes it difficult to draw firm conclusions from the results.

In a different approach, Experiment 3 acquired IVIM data at a much higher resolution in order to minimise the effects of partial voluming within voxels. Application of a rs-EPI sequence on a high gradient performance Prisma scanner allowed for the acquisition of ten 1 mm isotropic slices in a feasible scan time. The resulting images (see figure 5) show that *f_v_* and *f_v_D*^*^ are elevated in voxels containing CSF (as expected), but that no contrast is visible between grey and white matter voxels. In addition, there appears to be some consistent correlation between *f_v_* and CSF fraction, but not with GM or WM fraction, as shown in figure 7. Together, this strongly suggests that *f_v_* is primarily sensitive to voxelwise CSF content and not true capillary blood volume.

The box plots in figure 8 confirm that *f_v_* and *f_v_D*^*^ are highest in the CSF compartment, and also show that the median *f_v_D*^*^ value is marginally higher in WM than GM. The same observations were made for all subjects. Regardless of units, this result is both unexpected and unphysiological for a perfusion contrast (56). It also suggests that the plausible GM/WM ‘perfusion’ ratio observed in Experiment 1 may in fact be an averaged (GM+CSF)/WM ratio.

One possible explanation for the unexpectedly high WM results is overfitting. Throughout this study, data have been analysed using a biexponential fitting procedure for all voxels, even when the simpler monoexponential fit appears to do an adequate job. This was intentional and in accordance with the recent literature (29,37). However, by fitting a biexponential function to a noisy monoexponential decay curve, large values for *f_v_* can be estimated despite a true underlying *f_v_* = 0. The anisotropy in WM leads to greater variance in signal across diffusion directions compared to GM (see figure 6), which can masquerade as a lower SNR and may hence lead to poorer fitting.

As previously mentioned, the idea that the biexponential form of the diffusion signal arises from CSF contamination is not new (31). However the fact that this biexponential behaviour was observed even in 100% pure CSF voxels, with resulting *f_v_* of around 0.4, suggests that IVIM may in fact be measuring CSF motion (either turbulent or related to the cardiac cycle). In larger voxels, the mixing of different tissue types may be an additional contributing factor.

It is difficult to balance the requirements for high spatial resolution (to reduce partial voluming effects), high SNR (to ensure reasonable fitting) and acceptable temporal resolution in order to carry out a functional experiment with IVIM. However, one group has recently reported observing changes in *f_v_D*^*^ with hypercapnia (37), visual stimulation (38) and at different phases during the cardiac cycle (59), and interpreted these as increases in classical perfusion. Although absolute signal magnitude is very difficult to interpret, relative changes in IVIM parameters may reflect underlying physiological modulation, somewhat analogous to the blood oxygen level dependent (BOLD) signal. However, the advantages of this method over the BOLD signal are not clear, and the impact of any concurrent changes in CSF volume or pulsatility would need to be considered (60).

There have been a number of studies demonstrating that IVIM can be used to generate clinically useful contrasts in both the abdomen (15,20) and the brain (33,35). For these situations, it may not be important whether the contrast is generated by excess fluid (for example surrounding a tumour) or by true perfusion, and in all probability IVIM will – and should – continue to be a useful tool for creating these qualitative images. However, based on the results presented in this investigation, caution should be exercised in the interpretation of such images, especially if attempting to look beyond basic contrast to draw physiological conclusions from the data. Within the brain in particular, IVIM parameters *f_v_* and *f_v_D*^*^ do not appear to be related strongly to blood volume or perfusion, but rather to CSF content.

## Acknowledgements

EPSRC, MRC, NIHR Oxford Biomedical Research Centre

